# Deciphering the gene-centric spatial dynamics across tissue and time with GeneSPOT

**DOI:** 10.64898/2026.09.24.753111

**Authors:** Qi Zou, Zhikang Wang, Sijie Li, Senlin Lin, Chuangyi Han, Yan Cui, Shengxi Li, Lin Wang, Daoliang Zhang, Wei Zhang, Bin-Zhi Qian, Rui Gao, Zhiyuan Yuan

**Author notes:** These authors contributed equally to this work (Qi Zou, Zhikang Wang, Sijie Li). Corresponding Author:Wei Zhang, Bin-Zhi Qian, Rui Gao, Zhiyuan Yuan.

## Abstract

Spatial omics technologies provide unprecedented opportunities to investigate how molecular programs are organized within tissues. However, existing analytical frameworks primarily focus on identifying cellular states and characterizing differentially expressed molecular features across these states. The relational spatial organization of molecular features and its dynamics across biological contexts therefore remain largely unexplored. Here, we present GeneSPOT, a gene-centric computational framework for characterizing the spatial relationships among genes and quantifying their reorganization across biological conditions. Although gene-centric in design, GeneSPOT generalizes to other molecular features. Across diverse applications, GeneSPOT resolved tissue structures, integrated paired and unpaired spatial multi-omics data, unveiled the spatial convergence of gene programs during embryonic development, and identified aging-associated spatial reorganization among genes whose expression did not change significantly with age. GeneSPOT thus complements conventional cell-centric analysis, providing a framework for deciphering how spatial relationships among molecular features are reorganized across biological conditions and over time.

## Introduction

Biological processes within tissues are orchestrated by spatially organized molecular programs^1, 2^. Advances in spatial omics technologies have enabled spatially resolved profiling of molecular features across tissues and organs^3-9^. However, the prevailing analytical paradigm remains cell-centric, characterizing cellular spatial behaviors to model cell states^10-22^, interactions^23-29^, and communities^21, 30-41^. While this cell-centric perspective is invaluable, it leaves unresolved how the relational spatial organization of molecular programs varies across biological contexts, including different biological conditions (e.g., disease versus control), omics layers (e.g., transcriptomics and metabolomics), and dynamic processes (e.g., development).

Addressing this gap motivates a shift in the unit of analysis from cells to molecular features. To make this concept concrete without limiting its generality, we describe it in the context of spatial transcriptomics, where the molecular features are genes. The resulting gene-centric perspective repositions genes from static markers to active participants in coordinated spatial programs that define tissue states and drive biological processes, and may offer insights beyond those provided by cell-centric methods^42, 43^. Several methods for identifying spatially variable genes (SVGs) have adopted a gene-centric perspective^44-48^. For example, SpatialDE^49^ and SPARK^50^ characterize gene-wise spatial variation using statistical models, whereas STMiner^51^ applies optimal transport (OT) to separate this spatial variation from tissue-level background bias. Although effective for identifying SVGs, these methods are not designed to characterize the relational spatial organization underlying the functional landscape of genes.

Therefore, the spatial biology community currently lacks a unified framework for enabling this perspective shift, posing significant methodological challenges. These include (i) constructing biologically meaningful measures of spatial relationships among genes by integrating expression levels with spatial localization; (ii) reconciling disparate spatial coordinate systems and molecular landscapes across omics layers; (iii) generating an analyzable embedding space in which genes are organized according to their relative spatial relationships; and (iv) quantifying the dynamics of these spatial relationships across biological contexts, even when expression levels show no significant change.

To address these challenges, we introduce **Gene SP**atial **O**ptimal **T**ransport (GeneSPOT), a computational framework that models the relational spatial organization of genes. Analogous to a constellation, GeneSPOT positions genes within a context-specific Gene Constellation Embedding Space (GCES) according to their relative spatial relationships. To construct each GCES, GeneSPOT first represents the tissue as a spatial-aware cell graph, with cells as nodes and spatial proximity encoded by edges. For each gene, its expression across cells is normalized into a probability distribution over the graph, representing its spatial expression pattern. It then applies structure-preserving OT, using pairwise graph geodesic distances as transport costs, to quantify the minimum effort required to reconfigure one gene’s spatial distribution into that of another while respecting the underlying tissue structure. GeneSPOT also represents user-defined spatial landmarks as synthetic distributions that serve as anatomical anchors, enabling targeted analysis of how gene programs are organized relative to specific anatomical domains. The resulting gene-to-gene and gene-to-landmark transport profiles are integrated through graph representation learning^52^ to form a continuous GCES, in which genes and landmarks are organized according to their relative spatial relationships. By comparing changes in pairwise proximities among genes and landmarks across context-specific GCESs, GeneSPOT quantifies the reorganization of their spatial relationships across disease states, developmental stages, and aging.

We demonstrate the broad applicability and analytical utility of GeneSPOT across diverse tissues, molecular modalities, and biological timescales. Across these scenarios, GeneSPOT (i) organized genes into a functional landscape to delineate biologically meaningful spatial programs and resolve subtle transitional patterns within tissue architecture; (ii) uncovered cross-modal spatial relationships between transcriptomic and metabolomic features in both paired and unpaired settings (measured in the same or different samples, respectively); (iii) elucidated complex spatiotemporal dynamics in embryonic development, including the spatial convergence of hepatocyte and hematopoietic programs, which was not revealed by existing methods; and (iv) identified age-associated reorganization of spatial relationships among genes that was missed by conventional differential expression analyses during mouse brain aging. Together, these demonstrations underscore that GeneSPOT enables a shift toward a gene-centric paradigm for modeling the relational spatial organization of genes and decoding its dynamics across biological contexts.

## Results

### Overview of GeneSPOT

GeneSPOT is a gene-centric computational framework for representing the relational spatial organization of genes and tracking its dynamics across biological contexts. Although we use gene-centric terminology to follow conventions in spatial transcriptomics, GeneSPOT is omics-agnostic and applies to other molecular features, including proteins and metabolites. For concision, we refer to measurement units (e.g., cells, spots and bins) as “cells” and molecular features as “genes”, except when discussing specific multi-omics data.

GeneSPOT accepts spatial omics data at multiple resolutions, including single-cell and spot-level data (Figure 1A). For each biological context, the input comprises spatial coordinates and a corresponding feature expression matrix. GeneSPOT uses these inputs to construct a continuous GCES that organizes genes and user-defined landmarks according to their relative spatial relationships (Figure 1B). Within each context-specific GCES, gene-to-landmark proximity relates gene programs to anatomical domains, whereas gene-to-gene proximity captures spatial coupling among genes. Together, these complementary measures define the relational spatial organization of genes within tissue. Within this unified framework, GeneSPOT (i) preserves intrinsic tissue structure while reconciling disparate spatial coordinate systems and molecular landscapes; (ii) quantifies functional coupling among genes; and (iii) tracks the dynamics of spatial relationships among genes across biological conditions.

**Figure 1:**
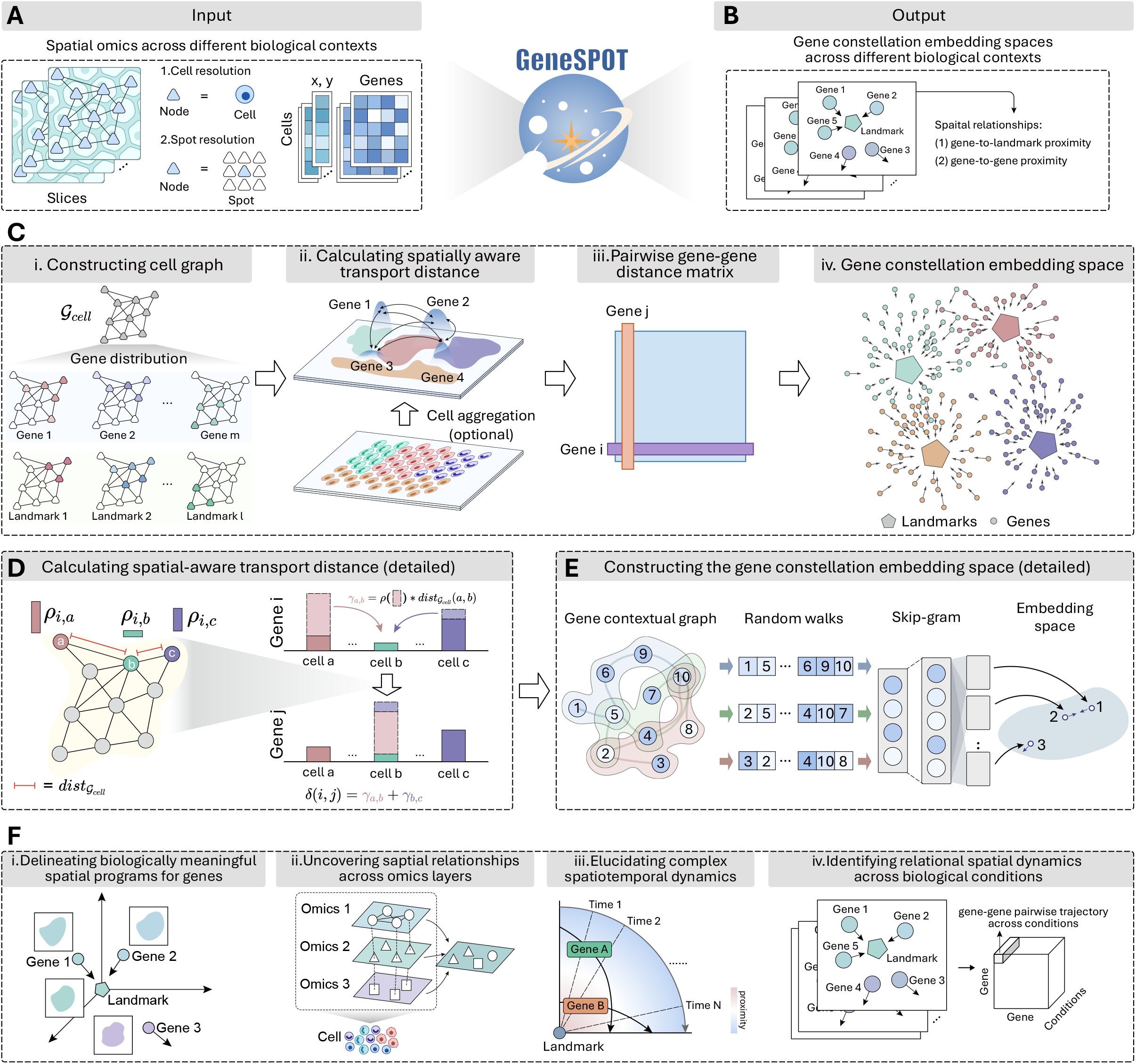
Conceptual overview of the GeneSPOT framework. GeneSPOT is a gene-centric computational framework for representing the relational spatial organization of genes and tracking its dynamics across biological contexts. **A**, GeneSPOT accepts spatial omics data at cell- or spot-level resolution and supports multiple analytical settings, including single-omics, paired multi-omics, unpaired multi-omics and spatiotemporal analyses. For each biological context, the input comprises spatial coordinates and a corresponding feature matrix (for example, gene expression). **B**, GeneSPOT generates a context-specific Gene Constellation Embedding Space (GCES), in which genes and spatial landmarks are positioned according to their relative spatial relationships. **C**, The GeneSPOT computational pipeline comprises four stages. (i) GeneSPOT constructs a spatial-aware cell graph that preserves the intrinsic tissue manifold and represents the spatial deployment of each gene as a probability distribution over this graph. User-defined spatial landmarks are incorporated as synthetic distributions restricted to user-specified regions of interest, providing biologically informed anchors for targeted analysis. (ii) Structure-preserving optimal transport is used to compute pairwise transport distances among genes and landmarks. (iii) The resulting pairwise distance profiles are used to construct a gene contextual graph, and biased random walks on this graph generate a topological corpus encoding spatial relationships among genes and landmarks. (iv) A skip-gram model uses this corpus to learn the final low-dimensional GCES, in which genes and landmarks with similar spatial deployment are positioned nearby. **D**, Detailed illustration of the spatial-aware transport distance calculation. Graph geodesic distances between cells define the transport cost, and the resulting transport distance quantifies the minimum effort required to reconfigure one spatial distribution into another while respecting the intrinsic tissue manifold. **E**, Detailed illustration of GCES construction using graph representation learning. The pairwise transport-distance profiles define a gene contextual graph. Biased random walks generate a topological corpus, which is then used to train a skip-gram model and learn the low-dimensional embedding. **F**, Key applications of GeneSPOT demonstrated in this work: (i) delineating biologically meaningful spatial gene programs, (ii) uncovering cross-modal spatial relationships, (iii) elucidating complex spatiotemporal dynamics, and (iv) identifying aging-associated spatial reorganization among genes whose expression did not change significantly with age.

The workflow for constructing each context-specific GCES comprises four stages (Figure 1C; see Methods). First, GeneSPOT constructs a spatial-aware cell graph that preserves the intrinsic tissue manifold and represents each gene as a probability distribution on the graph (Figure 1C i). GeneSPOT also incorporates user-defined spatial landmarks as synthetic distributions restricted to user-specified regions of interest, thereby supporting targeted biological inquiry (Figure 1C i). These landmarks serve as biologically informed anchors, providing a consistent basis for comparison across different biological contexts (see Methods). Second, GeneSPOT applies structure-preserving OT^43, 53-56^ to compute pairwise transport distances among genes and landmarks (Figure 1C ii). Specifically, the graph geodesic distances between cells define the corresponding transport cost (Figure 1D). The resulting pairwise transport distance therefore quantifies differences in spatial deployment by measuring the minimum effort required to reconfigure one distribution into another while respecting the intrinsic tissue manifold (Figure 1D; see Methods). Third, GeneSPOT uses the resulting pairwise distance profiles to construct a gene contextual graph and applies biased random walks to generate a topological corpus encoding spatial relationships among genes and landmarks (Figure 1C iii). Fourth, a skip-gram model uses this corpus to learn a low-dimensional GCES in which genes and landmarks with similar spatial deployment are positioned nearby (Figure 1C iv and Figure 1E; see Methods). Finally, GeneSPOT analyzes and compares context-specific GCESs across four analytical settings (Figure 1F; see Methods). These analyses (i) delineate biologically meaningful spatial gene programs; (ii) uncover spatial relationships across omics layers; (iii) elucidate complex spatiotemporal dynamics; and (iv) identify the reorganization of spatial relationships across biological conditions. For a comprehensive description of the GeneSPOT methodology, please refer to the Methods section.

### GeneSPOT Captures Biologically Meaningful Gene Programs

To assess GeneSPOT’s ability to resolve spatial gene programs in a highly organized tissue, we applied it to a public spatial transcriptomics dataset comprising three adult mouse medial prefrontal cortex (mPFC) slices^57^ (Figure 2A and Supplementary Table 1). We used BZ9 and BZ14 to construct a GCES containing four landmarks representing Layers I, II/III, V and VI, and reserved BZ5 as a held-out test slice (Figure 2B). Anatomical domain labels followed the original annotations^57, 58^ (Supplementary Figure 5; see Supplementary Notes 1.1).

**Figure 2:**
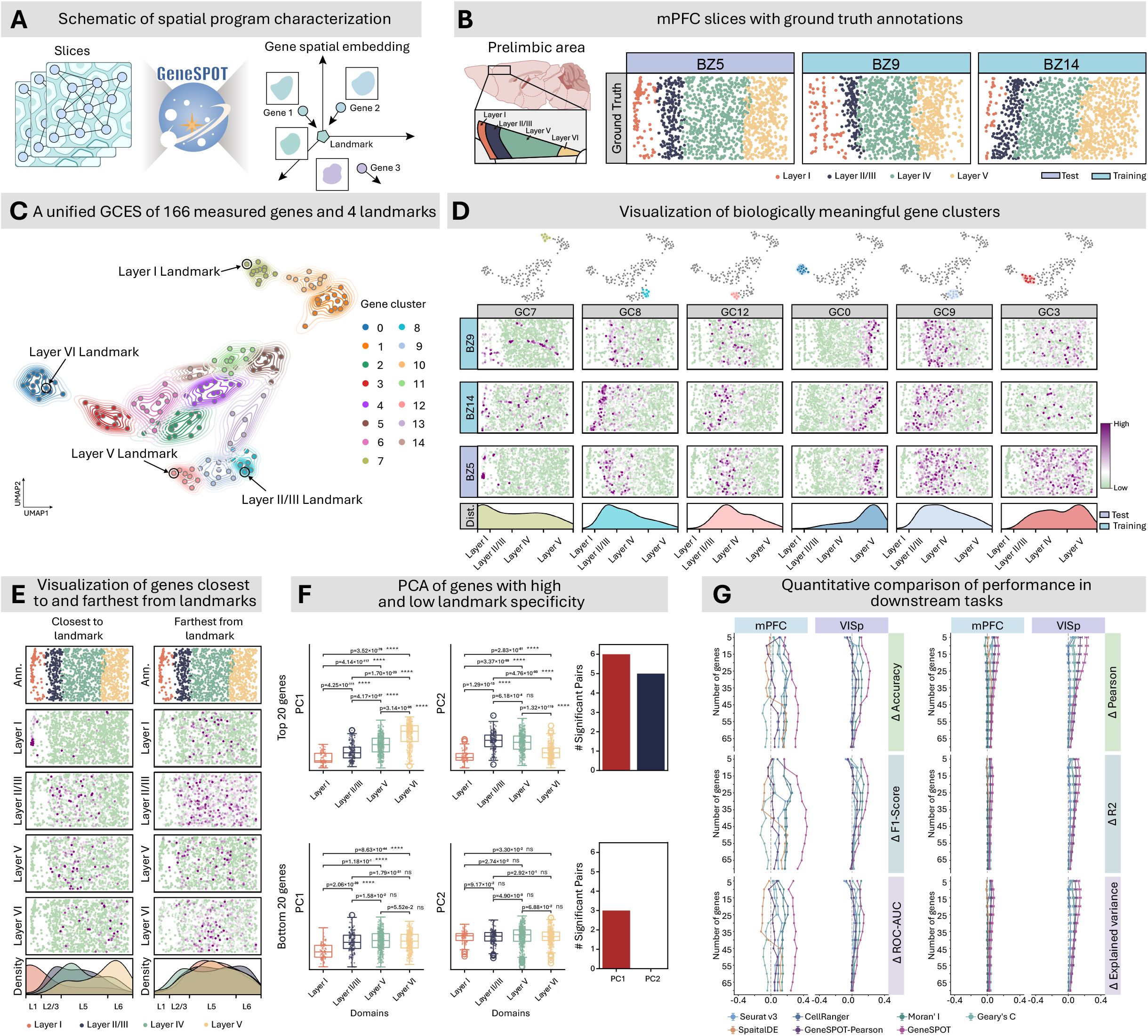
GeneSPOT captures biologically meaningful spatial gene programs in the mouse brain. **A**, Schematic diagram of the experimental setup. GeneSPOT was applied to generate a unified GCES that captures biologically informative gene programs. **B**, Spatial transcriptomics dataset comprising three adult mouse medial prefrontal cortex (mPFC) slices (BZ5, BZ9 and BZ14) with anatomical annotations for Layers I, II/III, V and VI. BZ9 and BZ14 were used to construct the GCES, whereas BZ5 was reserved as a held-out test slice. **C**, UMAP visualization of the unified GCES, in which 166 genes and four spatial landmarks corresponding to the cortical layers are co-embedded. Each point represents a gene or landmark, colored by its assigned gene cluster (GC). **D**, Visualization of representative GCs. The top row shows the position of each GC in the GCES. The next three rows show the average spatial expression of the genes in each GC in BZ9, BZ14 and BZ5, respectively. The bottom row shows the corresponding average expression profiles along the dorsal–ventral axis (x-axis), smoothed using kernel density estimation (y-axis). **E**, Spatial expression of genes closest to or farthest from each landmark in the held-out BZ5 slice. The top row shows the cortical layer annotations. The next four rows show the spatial expression of the gene closest to or farthest from each of the four landmarks. The bottom row shows the corresponding expression profiles along the dorsal–ventral axis (x-axis), smoothed using kernel density estimation (y-axis). Genes closest to a landmark exhibited sharply localized expression peaks within the corresponding anatomical domain, whereas genes farthest from the landmark lacked this localized expression. **F**, Unsupervised principal component analysis (PCA) of all 1,049 cells from the BZ5 slice (Layer I, n = 86 cells; Layer II/III, n = 167 cells; Layer V, n = 449 cells; Layer VI, n = 347 cells), using the top 20 genes with the highest standard deviation (STD) of their proximities across the four landmarks (top row) or the bottom 20 genes with the lowest STD (bottom row). The first two columns show the distributions of PC1 and PC2 scores across the four anatomical domains using boxplots (showing the median (center line), the 25th and 75th percentiles (box bounds), and whiskers extending to the most extreme observations within 1.5 times the interquartile range from the interquartile quartiles). Overlaid points represent individual cells. The third column shows the number of significantly separated domain pairs for each principal component. Pairwise comparisons were performed using two-sided unpaired Student’s t-tests with nominal *P* values and no adjustment for multiple comparisons (exact *P* values are shown above the comparisons). \*\*\*\**P* < 0.0001; ns, not significant. **G**, Quantitative benchmarking of gene prioritization performance in two supervised tasks using the mPFC and mouse visual cortex (VISp) datasets: a supervised domain classification task (left) and a supervised gene expression regression task (right). Performance was measured as the difference (Δ) in metric values between the top- and bottom-ranked gene sets across the indicated gene-set sizes. Error bars represent the s.e.m.

The resulting GCES positioned genes and spatial landmarks in a biologically meaningful configuration, with nearby points reflecting similar spatial deployment. In Figure 2C, each point represents a gene or landmark and is colored by its assigned gene cluster (GC). The resulting cluster organization aligned with known anatomical domains. Spatial visualization of the constituent genes, together with quantitative analyses, further confirmed that genes within these GCs were spatially enriched in the corresponding anatomical domains (Figure 2D, Supplementary Figure 1A-B). For example, GC 7 contained the Layer I landmark together with genes spatially enriched in that layer. Similarly, the constituent genes of GCs 8, 12 and 0 were enriched in Layers II/III, V and VI, respectively. Together, these results demonstrate that GeneSPOT organized genes according to their relative spatial relationships.

Beyond delineating domain-restricted gene clusters, the continuous GCES also captured nuanced transitional gene programs. Such programs can be missed by differentially expressed gene (DEG) and spatially variable gene (SVG) analyses because these approaches do not model relative spatial relationships among genes (Supplementary Figure 6). GeneSPOT instead identified numerous transitional programs spanning adjacent anatomical domains. Specifically, GC 9 was positioned between GC 8, associated with Layers II/III, and GC 12, associated with Layer V. The average spatial expression of its constituent genes accordingly spanned Layers II/III and V (Figure 2D and Supplementary Figure 1C-D). Functional enrichment analysis further showed that GC 9 was enriched for processes involved in inter-layer integration^59^ (Supplementary Figure 7). Similarly, GC 3 was positioned between GC 12 and GC 0, associated with Layers V and VI, respectively. Its constituent genes were expressed across both layers (Figure 2D and Supplementary Figure 1C-D). Together, these findings demonstrate that GeneSPOT resolves nuanced transitional programs that are not captured by standard DEG and SVG analyses.

To evaluate whether the constructed GCES reflected tissue organization, we examined spatial expression patterns in relation to gene-to-landmark proximity. This analysis showed that genes closest to a landmark exhibited sharply localized expression peaks within the corresponding anatomical domain (Figure 2E, left), whereas genes distant from the landmark lacked this localized expression (Figure 2E, right). To quantify this biological relevance, we ranked genes by the standard deviation of their proximities across the four landmarks. The top and bottom 20 genes represented the most and least landmark-specific spatial profiles, respectively. Using each gene set, we performed principal component analysis (PCA) on the held-out BZ5 slice and compared component scores between all pairs of anatomical domains. Domain-separation performance was defined as the number of domain pairs separated by two-sided unpaired Student’s t-tests at a nominal *P* < 0.05 (see Methods). On PC1, the top 20 gene set separated all six domain pairs, whereas the bottom 20 set separated three (Figure 2F and Supplementary Figure 8A-B). Across gene-set sizes ranging from 5 to 70, the top gene sets prioritized by GeneSPOT consistently outperformed competing methods and achieved complete pairwise separation with as few as five genes (Supplementary Figure 9). By contrast, GeneSPOT-Perm, in which anatomical domain labels were randomly permuted, failed to distinguish the domains (Supplementary Figure 9). Together, these results show that gene-to-landmark proximity identifies genes that are informative of anatomical organization.

GeneSPOT constructs an analyzable embedding space that facilitates the selection of spatially informative gene sets through gene-to-landmark and gene-to-gene proximities. To quantify this ability to prioritize genes that are both discriminative of anatomical domains and representative of the broader transcriptional landscape, we selected gene sets and trained models on the training slices, then evaluated them in two supervised tasks on a held-out test slice (Figure 2B; see Methods). First, a domain classification task tested whether a prioritized gene set could assign cells to their annotated anatomical domains. Strong performance in this task indicates that these genes are spatially organized relative to anatomical domains. Second, a leave-one-gene-out expression-regression task tested whether the expression of each gene could be predicted from that of its most proximal genes. Strong performance in this task indicates that the prioritized genes capture spatial coupling patterns among genes. An effective gene prioritization strategy should therefore yield strong performance for top ranked genes and weak performance for bottom ranked genes. We benchmarked GeneSPOT against highly variable gene (HVG) identification methods (Seurat^60^ and CellRanger^61^) and SVG identification methods (Moran’s I^62^, Geary’s C^62^, and SpatialDE^49^). We also included GeneSPOT-Pearson, an ablation of GeneSPOT in which Pearson correlation replaced structure-preserving OT. For a fair comparison, each baseline ranked genes using its native statistic (e.g., standardized variance for HVGs, spatial autocorrelation scores for SVGs; see Methods), and prioritization performance was defined as the difference between the top *k* and bottom *k* gene sets (see Methods). Across the evaluated values of *k*, GeneSPOT produced the largest top-to-bottom performance gaps in both domain classification and gene expression regression (Figure 2G). As shown in Supplementary Figure 10, its top ranked gene sets remained competitive with those prioritized by baseline methods, whereas its bottom ranked gene sets showed the lowest performance among all methods. Similar results were obtained in independent datasets from the mouse visual cortex^63^ (Figure 2G) and the mouse hypothalamus profiled by MERFISH^64^ (Supplementary Figure 2 and Supplementary Figure 3). Together, these results show that GeneSPOT prioritizes compact gene sets that are both spatially discriminative and representative of broader transcriptional programs.

To evaluate GeneSPOT’s ability to resolve fine-scale tissue architecture while retaining scalability and interpretability, we applied it to a high-resolution Xenium human breast cancer dataset (HBC) containing more than 160,000 cells^8^ (Supplementary Table 1). The results showed that GeneSPOT resolved the complex ductal epithelial architecture and revealed a continuous tumor-progression axis (Supplementary Figure 11–16; see Supplementary Notes 2). It also processed the dataset more than 260-fold faster than the comparator gene-centric method^51^ (Supplementary Table 2– 5). Detailed benchmarking results and analyses are provided in Supplementary Notes 2.

### GeneSPOT Integrates Paired Spatial Transcriptomics and Metabolomics Data to Uncover Cross-omics Spatial Relationships

One challenge in spatial omics is deciphering relationships between molecular features across omics layers, such as genes and metabolites, to achieve a holistic understanding of tissue organization^4, 65, 66^. GeneSPOT enables such analysis in paired multi-omics settings by linking features measured on the same tissue section and representing their relative spatial organization (Figure 3A and Figure 3B). To demonstrate this capability, we applied GeneSPOT to a paired transcriptomic and metabolomic dataset from a Parkinson’s disease (PD) mouse model^67^. Spatial transcriptomics (Visium) and metabolomics (MALDI mass spectrometry imaging, MALDI-MSI) were measured on the same brain slices (Figure 3A, left). Each slice contained an intact hemisphere and a hemisphere lesioned with 6-hydroxydopamine (6-OHDA) to model PD pathology (Figure 3A, right). We used one paired slice to construct the GCES and reserved another as a held-out test slice.

**Figure 3:**
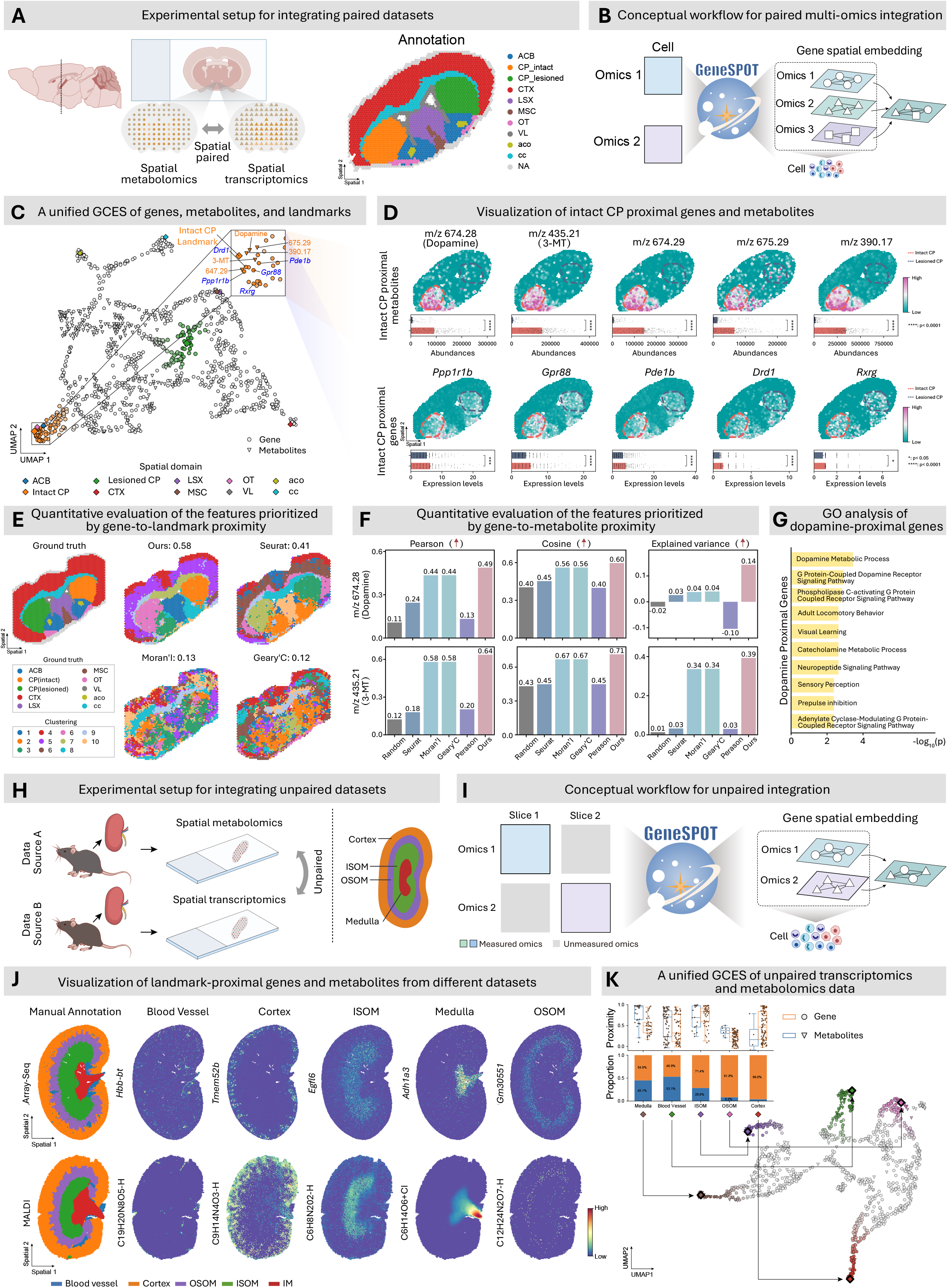
GeneSPOT integrates paired and unpaired spatial multi-omics data to uncover cross-omics spatial relationships. **A-G**, Integration of paired spatial transcriptomics and metabolomics from a Parkinson’s disease mouse model. **A**, Experimental setup for paired multi-omics integration. Spatial transcriptomics and metabolomics were measured on the same brain slices, each containing an intact hemisphere and a hemisphere lesioned with 6-hydroxydopamine (6-OHDA). One paired slice was used to construct the GCES, and another was reserved as a held-out test slice. B, Computational workflow for paired multi-omics integration. GeneSPOT co-embeds molecular features measured in different omics layers into a unified GCES. **C**, UMAP visualization of the unified GCES, in which genes (circles), metabolites (triangles) and spatial landmarks (diamonds) are co-embedded according to their relative spatial deployment. The inset highlights molecular features proximal to the intact choroid plexus (intact CP) landmark. **D**, Spatial distributions of intact-CP-proximal metabolites (top) and genes (bottom) identified by GeneSPOT. Color intensity indicates relative metabolite abundance or gene expression. Dashed contours indicate the regions used for quantitative analysis: red outlines denote the intact CP and correspond to the red bars, whereas blue outlines denote the lesioned CP and correspond to the blue bars. Bar plots show the mean ± s.e.m. abundance or expression of each feature across spots in the intact CP (n = 314 spots) and lesioned CP (n = 296 spots). Statistical significance was assessed using two-sided unpaired Student’s t-tests with nominal *P* values (exact *P* values were as follows: m/z 674.28: 4.19 × 10^−86^, m/z 435.21: 3.61 × 10^−153^, m/z 674.29: 1.06 × 10^−152^, m/z 675.29: 2.65 × 10^−52^, m/z 390.17: 6.23 × 10^−176^, *Ppp1r1b*: 8.38 × 10^−4^, *Gpr88*: 9.12 × 10^−7^, *Pde1b*: 2.41 × 10^−5^, *Drd1*: 2.3 × 10^−5^, *Rxrg*: 1.11 × 10^−2^). \*\*\*\**P* < 0.0001; \*\*\**P* < 0.001; \**P* < 0.05. **E**, Quantitative evaluation of features prioritized by gene-to-landmark proximity. Molecular features were prioritized on the training slice according to the standard deviation of their proximities to spatial landmarks and subsequently used to identify spatial domains in the held-out test slice. Concordance with the reference anatomical annotations was quantified using the adjusted Rand index (ARI). **F**, Quantitative evaluation of the features prioritized by gene-to-metabolite proximity. Linear regression models were trained on the training slice using gene sets prioritized separately by GeneSPOT, six comparison methods, and the GeneSPOT-coords ablation, and were evaluated on the held-out test slice. The models predicted the spatial abundance of dopamine (top row) and 3-methoxytyramine (3-MT; bottom row) across the indicated gene-set sizes. Performance was evaluated using Pearson correlation, cosine similarity and explained variance. **G**, Gene ontology analysis of dopamine-proximal genes revealed enrichment for relevant biological pathways (one-sided Fisher’s exact tests with Benjamini–Hochberg correction). **H-K**, Integration of unpaired spatial transcriptomics and metabolomics data from independent mouse kidney datasets. **H**, Experimental setup for unpaired multi-omics integration. **I**, Computational workflow for unpaired data integration using shared anatomical landmarks as conserved anchors across omics layers. **J**, Spatial expression and abundance maps of representative landmark-proximal genes and metabolites, respectively. **K**, UMAP visualization of the unified GCES constructed from the unpaired kidney datasets. Within this space, genes (circles), metabolites (triangles) and spatial landmarks (squares) are organized according to their inferred spatial relationships. Boxplots compare feature-to-landmark proximity distributions between genes and metabolites. Center lines indicate medians, box limits indicate the 25th and 75th percentiles, and whiskers extend to the most extreme observations within 1.5 times the interquartile range from the lower and upper quartiles. Overlaid points represent individual molecular features. The stacked bars show the proportions of genes and metabolites among the features proximal to each landmark.

The resulting GCES co-embedded genes (circles), metabolites (triangles), and spatial landmarks (diamonds), with proximity reflecting similarity in their spatial deployment (Figure 3C). This integration revealed biologically meaningful multi-omics programs associated with specific anatomical domains. For example, the intact choroid plexus (intact CP) landmark was positioned near the metabolites dopamine and 3-methoxytyramine (3-MT) and the genes *Pde1b*^68^, *Drd1*^69^, and *Gpr88*^70^ (Figure 3C). Spatial visualization and quantitative comparison across domains confirmed that these features were enriched in the intact CP domain (Figure 3D).

To quantitatively evaluate whether the integrated GCES captured tissue organization, we prioritized molecular features based on the standard deviation of their proximities to spatial landmarks. The prioritized features were then used for spatial clustering of the held-out test slice (see Methods). The features prioritized by GeneSPOT yielded greater concordance with the reference anatomical annotations (adjusted Rand index, ARI = 0.58) than features prioritized by Seurat (ARI = 0.41), Moran’s I (ARI = 0.13) or Geary’s C (ARI = 0.12) (Figure 3E). To determine whether this performance depended on the selection threshold, we varied the number of prioritized metabolites from 10 to 100 and the number of prioritized genes from 50 to 250 (see Methods). Across these ranges, feature sets prioritized by GeneSPOT consistently yielded higher ARI values than those prioritized by the baseline methods (Supplementary Figure 19). These results indicate that GeneSPOT-prioritized molecular features were more informative about tissue spatial organization than those prioritized by competing methods. Similar gene-prioritization performance was observed in an independent 10x Genomics Xenium breast cancer dataset^8^ (Supplementary Figure 4).

We next investigated whether the spatial distributions of metabolites could be predicted from the expression of their spatially coupled genes. To evaluate this capability, we defined a supervised prediction task using dopamine and 3-MT as target metabolites. Linear regression models were trained on the training slice to predict the spatial abundance of each metabolite from genes prioritized by different methods (see Methods). We evaluated predictive performance on the held-out test slice across different gene-set sizes (see Methods). Comparisons included four baselines (Random, Seurat, Moran’s I and Geary’s C) and two correlation-based rankings: Pearson (domain-level), which quantified correlations between features across anatomical domains, and Pearson (spot-level), which quantified raw abundance correlations across paired spots. The results showed that models trained with GeneSPOT-prioritized genes achieved the highest predictive performance across the evaluated settings, including higher performance than models based on spot-level Pearson rankings (Figure 3F). Whereas spot-level Pearson depends on local co-abundance and can be affected by sparsity and dropout, GeneSPOT models tissue-wide spatial deployment patterns of features, enabling it to capture spatial coupling even when local co-abundance is imperfect. To assess the contribution of the transport geometry, we also evaluated GeneSPOT-coords, an ablation of GeneSPOT that used pairwise Euclidean distances between cells as transport costs (see Methods). Models trained with genes prioritized by GeneSPOT outperformed those using genes prioritized by GeneSPOT-coords, supporting the value of structure-preserving transport geometry for capturing the intrinsic tissue manifold beyond Euclidean proximity (Figure 3F). Gene ontology analysis of the dopamine-proximal genes prioritized by GeneSPOT revealed enrichment for dopamine metabolism and signaling, as well as the catecholamine metabolic process (Figure 3G). These enrichments further support the biological coherence and functional relevance of the inferred multi-omics module. Together, these analyses show that GeneSPOT integrates paired spatial omics data into a unified representation that reflects anatomical organization and captures biologically coherent relationships across molecular layers.

### GeneSPOT Facilitates Integration of Unpaired Spatial Multi-Omics Datasets

Extending spatial multi-omics integration to unpaired datasets presents an additional challenge, as different molecular layers are profiled on separate tissue sections and may originate from different laboratories or large-scale consortia ^71-74^. Unlike paired measurements, unpaired datasets lack direct correspondence between measurement locations and therefore require a computational strategy capable of inferring cross-omics relationships through shared anatomical structures^75^. To facilitate the integration of unpaired multi-omics datasets, GeneSPOT uses shared spatial landmarks corresponding to matched anatomical domains as common anchors across omics layers (Figure 3H and Figure 3I).

To evaluate this capability, we integrated two independent mouse kidney datasets, one comprising spatially resolved transcriptomics^76^ and another providing spatially resolved metabolomics^77^ (Figure 3H). Although both datasets profiled the mouse kidney and shared underlying anatomical structures, such as the cortex and medulla, they originated from different studies and lacked cell-to-cell correspondence, providing a representative unpaired multi-omics integration setting (Figure 3I). Using these anatomical anchors, GeneSPOT co-embedded genes (circles), metabolites (triangles), and spatial landmarks (squares) into a unified GCES according to their relative proximities to the shared landmarks (Figure 3J and Figure 3K). Despite being measured in unpaired datasets, genes and metabolites proximal to each landmark in the GCES were spatially localized within the corresponding anatomical domain (Figure 3J). For example, GeneSPOT identified an inferred cross-omics association between *Ppp1r1b*, which encodes a signaling regulator, and a metabolite feature assigned the molecular formula C_13_H_9_ClO^−^, both of which were localized to the inner stripe of the outer medulla (ISOM). This spatial association suggests a potential link between *Ppp1r1b* associated signaling and local metabolite regulation, although the underlying mechanism remains to be determined.

The integrated GCES also revealed differences in the omics-layer composition of features proximal to different anatomical landmarks (Figure 3K). The blood-vessel landmark was associated with a relatively balanced set of genes and metabolites, whereas genes predominated among the features proximal to the medulla, ISOM, OSOM and cortex landmarks, particularly OSOM and cortex (Figure 3K, top left). Together, these results show that GeneSPOT links unpaired spatial omics datasets through shared anatomical landmarks and reveals candidate region-specific cross-omics relationships.

### GeneSPOT Quantifies Relational Shifts in Gene Programs Across Biological Contexts

Understanding how gene programs reorganize across biological contexts is important for characterizing processes such as disease progression^78^, tissue development^79^, and aging^80^. While a gene’s sequence defines its molecular identity, its functional role may vary with its relational spatial context, defined here by its spatial relationships with other genes^81^. Many existing cell-centric analyses, including those incorporating spatial information, primarily focus on changes in expression levels^80, 82, 83^ or cell state mapping^54, 84-86^, rather than on how relationships among genes reorganize across biological contexts.

GeneSPOT quantifies these dynamics by comparing proximities across context-specific GCESs. Specifically, changes in gene-to-landmark proximities reveal shifts in the anatomical anchoring of genes, whereas changes in gene-to-gene proximities characterize the reorganization of spatial coupling among genes (see Methods). Together, these complementary measures provide a relational account of functional shifts and track how gene programs spatially reorganize during biological processes. We applied this framework to characterize spatiotemporal dynamics during mouse embryonic development^79^ (Figure 4) and age-dependent reorganization of spatial relationships between genes in the aging mouse brain^80^ (Figure 5).

**Figure 4:**
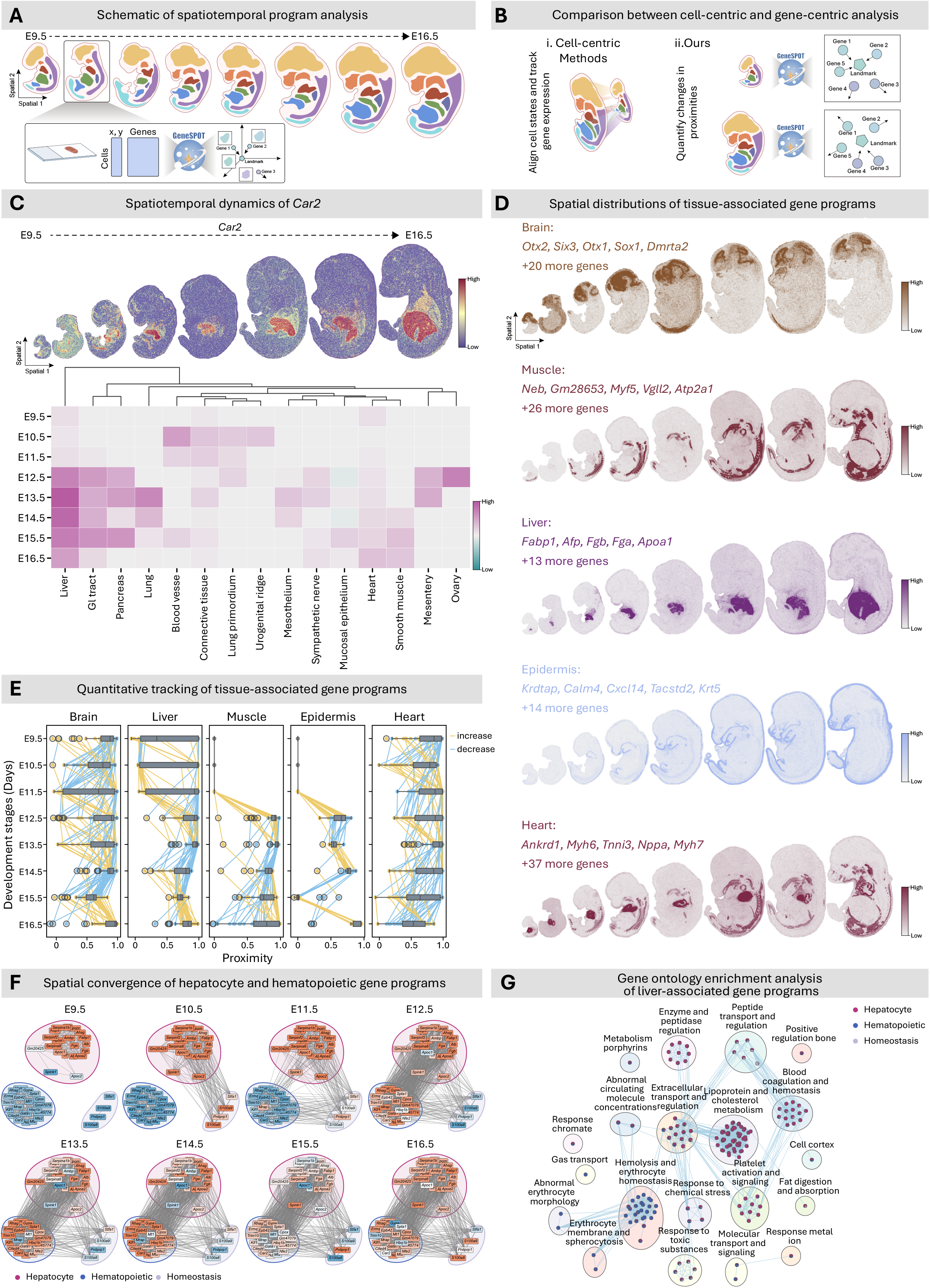
GeneSPOT unveils spatiotemporal gene program dynamics during mouse embryogenesis. **A**, Schematic of the spatiotemporal gene-program analysis. GeneSPOT was applied to a Stereo-seq atlas of mouse organogenesis spanning eight developmental stages from E9.5 to E16.5. Stage-specific GCESs were constructed using annotated tissue structures as spatial landmarks, yielding a spatiotemporal atlas of gene program dynamics. **B**, Comparison of cell-centric and gene-centric analyses. Cell-centric approaches align cell states or track gene expression across developmental stages, whereas GeneSPOT quantifies changes in gene-to-landmark and gene-to-gene proximities. **C**, Spatiotemporal dynamics of the gene *Car2*. The heatmap shows its proximity to tissue landmarks across developmental time, with rows representing developmental stages and columns representing tissue landmarks. *Car2* shows diffuse associations with early organ primordia at E9.5–E10.5, followed by increasingly specific associations with developing organs, including the liver and heart. **D**, Spatial distributions of tissue-associated gene programs defined by genes proximal to the brain, muscle, liver, epidermis and heart landmarks across the indicated developmental stages. **E**, Gene-to-landmark proximity trajectories for genes associated with the brain, liver, muscle, epidermis and heart landmarks. Each line represents one gene, and yellow and blue segments indicate increases and decreases, respectively, in proximity between consecutive developmental stages. Points represent individual genes (brain, n = 25 genes; liver, n = 18 genes; muscle, n = 31 genes; epidermis, n = 19 genes; heart, n = 42 genes). Center lines indicate medians, box limits indicate the 25th and 75th percentiles, and whiskers extend to the most extreme observations within 1.5 times the interquartile range from the lower and upper quartiles. **F**, Dynamic convergence of hepatocyte and hematopoietic gene programs. Networks show genes with proximity to the liver landmark greater than 0.45 at each developmental stage. Node colors encode gene-to-liver-landmark proximity at the corresponding developmental stage. Edges connect gene pairs with gene-to-gene proximity greater than 0.5, and edge width scales with gene-to-gene proximity. **G**, Pathway enrichment analysis supports the distinct functional identities of the hepatocyte- and hematopoietic-proximal gene modules. In the enrichment graphs, nodes represent gene ontology terms, and edges connect terms with overlapping genes. The pie chart within each node shows the component distribution of the hepatocyte (red), hematopoietic (blue), and homeostasis (purple) modules for each enriched gene ontology term, revealing functionally related clusters such as lipid metabolism for hepatocytes and erythrocyte differentiation for hematopoietic programs.

**Figure 5:**
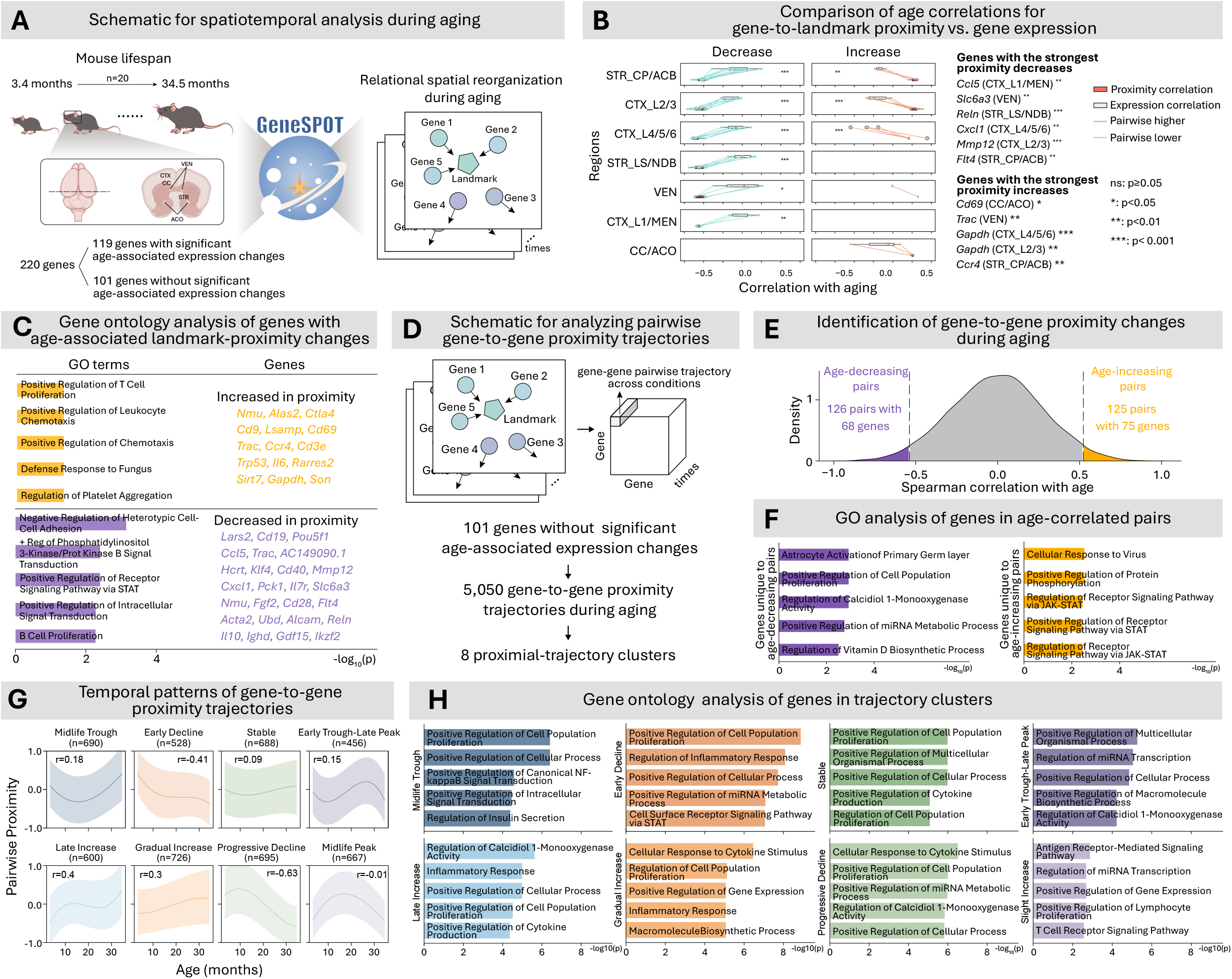
GeneSPOT discovers age-dependent spatial reorganization among genes during mouse brain aging. **A**, Schematic of the analysis of a mouse brain aging dataset spanning the entire lifespan from 3.4 to 34.5 months. **B**, Comparison of Spearman correlations with age for gene-to-landmark proximity and gene expression across the indicated brain regions. Among genes without significant age-associated expression changes, “increased genes” were defined as those with a Spearman correlation between age and gene-to-landmark proximity greater than 0.3, a lower bound of the 95% confidence interval greater than 0 and a nominal two-sided Spearman *P* < 0.05. “Decreased genes” were defined as those with a Spearman correlation less than −0.3, an upper bound of the 95% confidence interval less than 0 and a nominal two-sided Spearman *P* < 0.05. Boxplot center lines indicate medians, box limits indicate the 25th and 75th percentiles, and whiskers extend to the most extreme observations within 1.5 times the interquartile range; lines connect the proximity- and expression-based age-correlation coefficients for the same gene. Genes with the strongest negative or positive age correlations in gene-to-landmark proximity are shown on the right. Exact nominal *P* values for the negatively correlated pairs were: *Ccl5* (CTX_L1/MEN): 5.16 × 10^−3^; *Slc6a3* (VEN): 6.75 × 10^−4^; *Reln* (STR_LS/NDB): 1.39 × 10^−3^; *Cxcl1* (CTX_L4/5/6): 4.2 × 10^−4^; *Mmp12* (CTX_L2/3): 1.7 × 10^−3^; *Flt4* (STR_CP/ACB): 5.16 × 10^−3^. Exact nominal *P* values for the positively correlated pairs were: *Cd69* (CC/ACO): 1.79 × 10^−2^; *Trac* (VEN): 7.67 × 10^−3^; *Gapdh* (CTX_L4/5/6): 6.5 × 10^−4^; *Gapdh* (CTX_L2/3): 2.2 × 10^−3^; *Ccr4* (STR_CP/ACB): 6.92 × 10^−3^. **C**, Gene ontology analysis of genes with age-associated changes in gene-to-landmark proximity (one-sided Fisher’s exact tests with Benjamini–Hochberg correction). **D**, Schematic of the analysis of pairwise gene-to-gene proximity trajectories. For 101 genes without significant age-associated expression changes, GeneSPOT quantified gene-to-gene proximity across the aging time course for each of the 5,050 unique gene pairs. **E**, Distribution of Spearman correlation coefficients between age and gene-to-gene proximity across the 5,050 gene-pair trajectories. Age-decreasing pairs were defined by correlations below the 2.5th percentile of the empirical distribution and a nominal two-sided Spearman *P* < 0.05, whereas age-increasing pairs were defined by correlations above the 97.5th percentile and a nominal two-sided Spearman *P* < 0.05. **F**, Gene ontology analysis of genes unique to the age-decreasing or age-increasing gene-pair sets (one-sided Fisher’s exact tests with Benjamini– Hochberg correction). **G**, Temporal patterns of all pairwise gene-to-gene proximity trajectories. Smoothed median and interquartile range (error band) of gene-to-gene proximity z-scores across ages are shown for all gene-pair trajectories assigned to each cluster; shaded bands indicate the interquartile range. Pearson correlation (r) between median gene-to-gene proximity z-score and age is indicated; the number of trajectories within each cluster is noted in parentheses. **H**, Gene ontology analysis of the genes within each trajectory cluster (one-sided Fisher’s exact tests with Benjamini–Hochberg correction).

### GeneSPOT Unveils the Spatiotemporal Dynamics of Gene Programs during Embryonic Development

To unveil the dynamics of gene programs during organogenesis, we applied GeneSPOT to a large-scale Stereo-seq dataset spanning eight stages from E9.5 to E16.5 of mouse embryonic development^79^ (Figure 4A). By constructing a GCES for each developmental stage using annotated tissue structures as landmarks, we created a spatiotemporal atlas of gene program dynamics (see Methods). Whereas cell-centric methods primarily infer cellular trajectories^54, 84-86^ or track gene expression levels^80, 82, 83^ across timepoints (Figure 4B, left), GeneSPOT quantifies the spatial reorganization of gene programs by comparing pairwise proximities across stage-specific GCESs (Figure 4B, right).

We first tracked the spatiotemporal organization of individual genes, using Car2 as an example. Its gene-to-landmark proximity showed diffuse associations with early organ primordia at E9.5-E10.5, followed by increasingly specific associations with mature organs such as the liver and heart (Figure 4C). This spatial consolidation was consistent with the known functional specialization of *Car2* in physiological processes during organ maturation^87-89^.

Beyond individual genes, GeneSPOT also revealed the dynamic landscape of genes associated with specific tissues across developmental stages. GeneSPOT identified and tracked the dynamics of landmark-proximal gene programs for major tissues, including brain, liver, muscle, epidermis, and heart, over developmental time (Figure 4D and Figure 4E). These landmark-proximal programs recapitulated pivotal morphogenetic transitions, such as the pronounced morphological expansion of the liver between E13.5 and E14.5^90^ (Figure 4D, Liver), and the structural stabilization of the epidermis during mid-to-late stages^91^ (Figure 4D, Epidermis). GeneSPOT also reveals a continuous spatial progression from mesenchymal compartments to differentiated muscle structures (Figure 4D, Muscle). Moreover, coordinated developmental patterns were observed between the heart and adjacent muscle tissues, suggesting shared spatial programs and synchronized maturation^92^ (Figure 4D, Heart). Together, these results show that GeneSPOT quantitatively captures the spatial dynamics of tissue-associated gene programs across developmental stages.

Such a dynamic landscape also reflects the complex events of organogenesis. GeneSPOT goes beyond simply identifying upregulated markers to quantifying the spatiotemporal synchronization of functional modules. A notable example was the spatial convergence of hepatocyte and hematopoietic gene programs (Figure 4F). While previous analyses viewed liver development as the parallel upregulation or downregulation of markers^90^, GeneSPOT revealed that these two modules, initially spatially disjoint, became increasingly coupled in relational space. This convergence was characterized by increasing cross-module connectivity between an apolipoprotein hub in the hepatocyte program and an erythrocyte-membrane hub in the hematopoietic program (Figure 4F). By identifying candidate bridge genes between these modules, GeneSPOT revealed a potential molecular interface between hepatocyte and hematopoietic programs. This inferred spatial convergence was consistent with the established role of the fetal liver as the primary site of hematopoiesis^93^. Pathway analysis supported the distinct functional identities of these modules, with hepatocyte-proximal genes enriched for lipid metabolism and hematopoietic-proximal genes enriched for erythrocyte differentiation (Figure 4G). This systems-level insight into the coordinated spatial convergence of entire functional modules provides a layer of understanding that is not readily apparent from traditional cell-centric analyses.

### GeneSPOT Discovers Age-associated Reorganization of Gene Spatial Relationships during Mouse Brain Aging

We next examined age-associated reorganization of gene spatial relationships using a single-cell spatial transcriptomics atlas of the aging mouse brain spanning 20 time points from 3.4 to 34.5 months^80^ (Figure 5A). We specifically investigated whether GeneSPOT could identify age-dependent changes in spatial relationships among genes without significant age-associated expression changes. Such relational changes may be missed by conventional expression-based analyses (Figure 5A; see Methods).

GeneSPOT identified numerous genes with age-associated changes in gene-to-landmark proximity, although their expression levels were not significantly correlated with age (Figure 5B; see Methods). Genes showing significant decreases in landmark proximity were enriched for pathways related to cellular homeostasis, signaling, and plasticity, consistent with weaker anatomical anchoring (Figure 5C). For instance, in the cortical layers (CTX_L1/MEN), *Ccl5, which encodes a chemokine*^*94*^, was among the genes with the strongest decreases in landmark proximity with age (Spearman correlation with age *ρ*=-0.60, two-sided *P=*5.16 × 10^−3^), while its expression showed only a weak correlation with age (Spearman correlation with age *ρ*=0.21, two-sided *P=*3.6 × 10^−1^) (Figure 5B). By contrast, genes showing significant increases in landmark proximity were enriched for immune activation and inflammatory-response pathways, consistent with stronger anatomical association (Figure 5C). One example is *Cd69*, which encodes a classical marker of T-cell tissue residency and activation involved in retaining immune cells within non-lymphoid tissues^*95, 96*^. Within the corpus callosum and anterior commissure (CC/ACO), *Cd69* expression was not significantly correlated with age (Spearman’s ρ=0.19, two-sided *P=*4.2 × 10^−1^). However, its proximity to the CC/ACO landmark increased significantly with age (Spearman’s ρ=0.52, two-sided *P=*1.79 × 10^−2^) (Figure 5B).

This contrast showed that gene-to-landmark proximity captured age-associated shifts in anatomical anchoring that were not apparent from expression levels alone.

Beyond changes in individual gene-to-landmark proximity, GeneSPOT captured the temporal reorganization of gene-to-gene spatial relationships by tracking gene-to-gene proximity trajectories over time. Among the 101 genes without significant age-associated expression changes, we computed pairwise gene-to-gene proximity trajectories across the aging time course (Figure 5D). This analysis revealed two modes of reorganization during aging (Figure 5E). One mode was spatial decoupling, in which gene-to-gene proximity decreased with age. Genes specific to decoupling pairs were enriched for astrocyte activation, cell population proliferation and vitamin D- and miRNA-related processes (Figure 5F). This pattern suggests that aging may disrupt the spatial coordination of astrocyte-associated, proliferative and regulatory gene programs. The other mode was spatial coupling, in which gene-to-gene proximity increased with age. Genes specific to coupling pairs were enriched for cellular responses to viruses, protein phosphorylation and JAK–STAT signaling (Figure 5F). This pattern suggests that inflammaging may involve the progressive spatial synchronization of antiviral- and JAK–STAT-associated gene programs. Together, these findings reveal how spatial relationships between genes are reorganized during brain aging, providing relational insights that were not apparent from expression levels alone.

To further determine whether these proximity trajectories reflected biological programs, we clustered all pairwise gene-to-gene proximity trajectories into eight temporal patterns: Midlife Trough, Early Decline, Stable, Early Trough–Late Peak, Late Increase, Gradual Increase, Progressive Decline, and Midlife Peak (Figure 5G). Functional annotation revealed both shared and pattern-specific biological associations across these trajectory clusters (Figure 5H). The Early Decline and Progressive Decline clusters were enriched for genes involved in inflammatory or cytokine responses, cell population proliferation, miRNA regulation, and vitamin D-related processes. Despite their opposing trajectory directions, the Late Increase and Gradual Increase clusters were also associated with cytokine and inflammatory responses, together with regulation of gene expression. The non-monotonic patterns captured additional functional associations: the Midlife Trough cluster was enriched for NF-κB-related and intracellular signaling processes, the Early Trough–Late Peak cluster for miRNA transcription and multicellular organismal processes, and the Midlife Peak cluster for antigen receptor and T-cell receptor signaling. The Stable cluster was associated primarily with proliferation- and cytokine-related processes. Together, these results show that genes involved in related biological processes can exhibit distinct temporal trajectories of spatial coupling, revealing heterogeneous spatial reorganization that was not apparent from expression-level changes alone.

## Discussion

In this study, we introduce GeneSPOT, a gene-centric computational framework for spatial omics analysis. By shifting the unit of analysis from cells to molecular features, GeneSPOT complements conventional cell-centric approaches and enables the relational spatial organization of molecular features and its dynamics to be investigated across biological contexts. Across diverse applications, GeneSPOT resolved complex tissue architecture, integrated paired and unpaired spatial multi-omics data, and quantified the reorganization of spatial gene programs across biological contexts. These analyses revealed the spatial convergence of functionally distinct gene programs during development and age-associated reorganization of spatial relationships among genes whose expression did not change significantly with age.

While GeneSPOT offers considerable strengths, its reliance on shared landmarks for integrating unpaired data poses a challenge when few reliably corresponding anatomical domains exist across datasets. Additionally, GeneSPOT employs a transductive learning approach, meaning that the model may require retraining when new molecular features, landmarks, or biological contexts are introduced. Developing an inductive version of GeneSPOT that generalizes to new contexts without full retraining would broaden its applicability^97^. GeneSPOT identifies associative changes in spatial organization; functional and mechanistic interpretations therefore require independent experimental validation.

Together, GeneSPOT provides a complementary framework for analyzing spatial omics data from the perspective of molecular features. As spatial omics resources continue to expand across molecular modalities and biological contexts, GeneSPOT may facilitate comparative analyses of how coordinated molecular programs are organized and reorganized across tissues and over time.

## Acknowledgements

The authors have no acknowledgements to declare.

## Author Contributions Statement

Z.Y. and R.G. conceived and designed the overall study. Z.Y., R.G., W.Z., and B.Q. supervised the project. Z.Y. and Q.Z. designed and developed the computational methods. Data collection and initial preprocessing of all datasets were conducted by Q.Z., Z.W., S. Lin, and Y.C. The primary analyses and figure generation were performed by Z.Y., Q.Z. and Z.W. L.W. and Sh. Li specifically contributed to the metabolic analysis by applying GeneSPOT to integrate spatial transcriptomics with metabolomics data. B.Q. provided key insights and guidance on the breast cancer data analysis. The manuscript was written by Z.Y. and Q.Z. with significant review and polishing from Z.W., Si. Li, C.H., D.Z., W.Z., and the supervisory team. All authors have read and approved the final version of the manuscript.

## Methods

GeneSPOT is a gene-centric computational framework for characterizing the relational spatial organization of genes and quantifying its dynamics across biological contexts. Although the term “gene-centric” aligns with established conventions in spatial transcriptomics, the framework is inherently omics-agnostic and generalizes to other molecular features, such as proteins and metabolites. At its core, GeneSPOT constructs a Gene Constellation Embedding Space (GCES) for each biological context (e.g., a tissue slice, developmental stage, or disease state). Together, these context-specific GCESs facilitate quantitative comparison of spatial gene programs across contexts. The pipeline consists of four main steps: (1) constructing a spatial-aware cell graph; (2) computing pairwise spatial transport distances among genes and landmarks using structure-preserving optimal transport (OT); (3) generating an analyzable GCES using graph representation learning; and (4) quantifying relational spatial dynamics by comparing gene-to-landmark and gene-to-gene proximities across context-specific GCESs. The first three steps are performed separately for each biological context, whereas the fourth compares relational spatial organization across contexts.

### Input

GeneSPOT accommodates spatial omics data at multiple resolutions, ranging from high-resolution cell-level measurements to low-resolution spot-level measurements. This framework supports diverse analytical scenarios, including: (1) single-omics analysis, where one omics layer (e.g., transcriptomics) is measured across all slices; (2) paired multi-omics analysis, where multiple omics layers (e.g., transcriptomics and metabolomics) are simultaneously measured on the same slice; and (3) the more complex unpaired multi-omics analysis, where different omics layers are measured on separate slices. For clarity, the units of measurement (e.g., cells, spots, bins) will be collectively referred to as “cells” for brevity.

The primary function of GeneSPOT is to generate a unique context-aware GCES for each specified biological condition. A biological context comprises one or more tissue slices analyzed jointly under a specified experimental condition, developmental stage, or disease state. Although GeneSPOT can accommodate diverse spatial omics scenarios (e.g., single-omics, paired multi-omics, unpaired multi-omics), we detail its core methodology using single-omics spatial transcriptomics. For brevity, all measured omics features are hereafter referred to as “genes”. Specifically, a given slice, comprising a set of cells 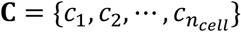, is characterized by their spatial coordinates 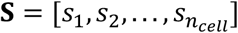 and gene expression matrix 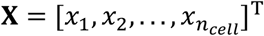. Here, *n*_*cell*_ indicates the total number of cells, 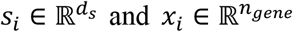 indicate the spatial coordinate and gene expression profile of *i*_*th*_ cell, respectively.

### Construction of the spatial-aware cell graph

In contrast to dissociated single-cell data, spatial omics data are characterized by the preservation of spatial context during profiling. This provides a distinct advantage for quantifying the in-situ organization and distribution of genes, i.e., spatial deployment. GeneSPOT achieves this by modeling spatial omics data as a spatial-aware cell graph. Here, each graph can be represented as *g*_*cell*_ = (C, *E*_*cell*_), where C denotes the cell nodes characterized by their gene expression profiles *X* and *E*_*cell*_ is a set of edges indicating the pairwise spatial-aware relationships.

GeneSPOT does not rely solely on raw physical coordinates. Although Euclidean distances are useful for applying spatial constraints, they are inadequate for multi-slice integration because physical coordinates do not correspond biologically across different tissue sections. Moreover, relying solely on physical proximity can merge regions that are spatially adjacent but biologically distinct (e.g., separated by an anatomical septum), resulting in transport through empty physical space rather than along the biological structure. To overcome this limitation, GeneSPOT reconstructs the intrinsic tissue manifold by embedding cells into a spatial-aware cell embedding space **H** that integrates both relative spatial proximity and gene expression profiles. Specifically, we first utilized the **S** and **X** as input for the state-of-the-art domain identification methods (e.g., MENDER^31^, STAGATE^34^). Given that spatial information as well as gene expression profiles were explicitly incorporated into the learning process, the resulting latent features are confirmed to be spatial-aware cell embeddings. Afterwards, the spatial-aware relationships are formally defined by constructing an undirected k-Nearest Neighbor (kNN) graph based on the spatial-aware cell representations **H**. An edge (*u, v*) is created if the Euclidean distance *dist*(*h*_*u*_, *h*_*v*_) between cell *c*_*u*_ and cell *c*_*v*_ is among the top *k*_spatial_ nearest neighbors of the other:

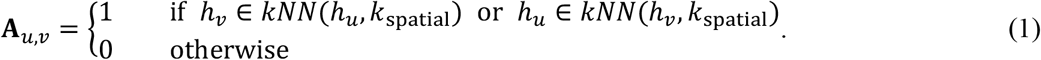

Here, *kNN*(*h*_*u*_, *k*_spatial_) denotes the set of *k*_spatial_ nearest neighbors of cell *c*_*u*_ based on Euclidean distances in the unified cell representation space **H**. The resulting spatial-aware cell graph *g*_*cell*_ delineates the discrete manifold structure of the cellular environment, serving as the transport geometry that preserves tissue structure that preserve tissue structure. Crucially, because this graph is defined by a joint spatial–expression representation rather than raw physical coordinates alone, it enables the seamless integration of cells across multiple slices.

### Calculate spatial-aware transport distance via structure-preserving optimal transport between genes

With the *g*_*cell*_ constructed, GeneSPOT conceptualizes the spatial deployment of each gene as a probability distribution of mass over the nodes of spatial graph *g*_*cell*_. The difference between the spatial deployment of any two genes is then formally quantified as the effort required to transport one gene’s distribution to match the other, constrained by the path defined by the graph’s edges. Specifically, the probability distribution ρ_*i*_ for a given gene *g*_*i*_ is defined by normalizing its expression profiles across all cells:

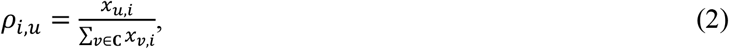

where *x*_*u,i*_ is the expression profile for *g*_*i*_ on cell node *c*_*u*_. We then quantify the spatial-aware transport distance between two spatial distributions ρ_*i*_ and ρ_*j*_ by the graph-based Wasserstein distance:

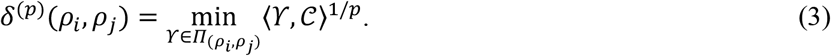

Here, *p* **∈** [1, *∞*) refer to the *p*-norm used when computing the Wasserstein distance, with a default value of 1. The cost matrix 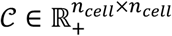 has entries 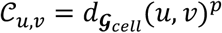 defined by the geodesic distance across 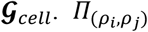 represents the set of feasible transport plans ϒ that push the mass from the source distribution ρ_*i*_ to the target distribution ρ_*j*_ where:

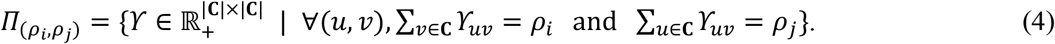

This design facilitates transport along the underlying biological structure rather than through empty physical space, thereby preserving the microstructure.

### Construct the gene constellation embedding space

To organize gene programs into a functional landscape, GeneSPOT uses the calculated spatial-aware transport distances to further construct an undirected gene contextual graph *g*_*gene*_ = (**G, E**_*gene*_). Here, *G* represents the gene nodes, and *E*_*gene*_ is a set of edges that connect each gene to its *k*_*gene*_ nearest neighbors as determined by the calculated transport distances. This gene contextual graph serves as a “document” that records the spatial relationships among genes. Following the gene contextual graph, we generate a topological corpus by performing biased random walks. The resulting sequence of visited nodes is treated as “sentences”, with the individual gene acting as “words”. Specifically, for each node *g*_i_ **∈** *G*, a “sentence” 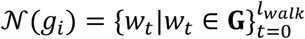 is generated by performing random walks starting from itself. Here, *l*_*walk*_ indicates the fixed walk length, *w*_*t*_ denotes the *t*-th node in the walk starting with *w*_0_ = *g*_*i*_. For *t ≥* 2, suppose that the walk arrives at the current node *u* = *w*_*t*−1_ from the previous node *s* = *w*_*t*−2_. The probability of transitioning to a candidate neighboring node *v* = *w*_*t*_ is defined as:

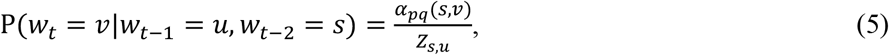

where *Z*_*s,u*_ is the normalizing constant overall all neighbors of *u*, and:

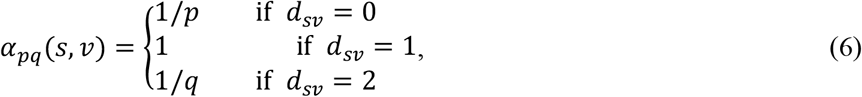

where *d*_*sv*_ represents the shortest path distance between the previous node *s* and the potential next node *v*. The parameter *p* and *q* guide the balance between local and global exploration of the graph. In this study, *p* and *q* are set as 1 and 0.5, respectively, by default.

The topological corpus is then fed into a skip-gram model to learn the low-dimensional gene embeddings, forming the GCES. The model trains a mapping function 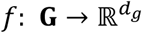 to map gene nodes with similar local context to nearby points in GCES. The training objective is to maximize the probability of predicting the local context for each node:

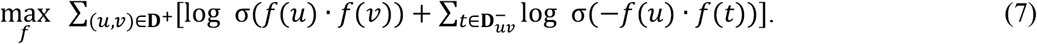

Here, **D**^*+*^ represents the set of positive samples, which are pairs of nodes (*u,v*) that co-occur within a defined context window size *k*_*context*_in the random walk sequence 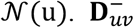 is the set of *k* _*negative*_ negative samples for each positive pair, which is drawn from the uniform distribution. In this study, we set *k*_*negative*_ = 1. σ(·) is the sigmoid function. Optimizing this objective function encourages similarity for true context pairs while simultaneously promoting dissimilarity for negatively sampled pairs. Consequently, proximity in the GCES reflects similarity in spatial deployment, and the overall structure of the GCES provides a low-dimensional, interpretable representation of relational spatial organization.

### Define spatial landmarks to facilitate biologically informed analysis

To facilitate biologically informed analysis, GeneSPOT integrates user-defined spatial landmarks as well as measured omics data for joint computation. Specifically, a landmark, *l* **∈ L**, is represented as a synthetic spatial distribution restricted to a user-defined region of interest (e.g., a user-defined spatial domain) corresponding to a subset of cells C_*l*_ *⊆* C. The landmark profile, 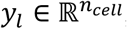, is formally represented as:

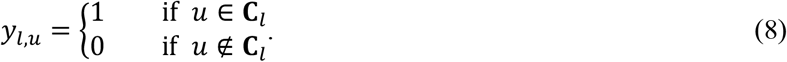

These landmark profiles are then treated as additional features and concatenated with the endogenous gene expression matrix *X*, resulting in an expanded matrix 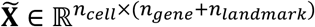. Here, *n*_*landmark*_ is the total number of landmarks. By jointly modeling genes and landmarks in 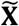, GeneSPOT quantifies the spatial relationships of genes relative to fixed biological reference landmarks, thereby anchoring the comparative analysis across disparate biological contexts, such as disease states and development stages.

### Cell aggregation for computational efficiency

To enhance the scalability of GeneSPOT in large-scale spatial omics datasets comprising millions of cells, we introduce a graph coarse-graining strategy. This approach aggregates cells in the spatial-aware graph *g*_*cell*_ into meta-cells based on spatial landmarks or user-defined biological annotations, representing each meta-cell by the cumulative profile of its constituent cells. Subsequently, GeneSPOT utilizes the Partition-Based Graph Abstraction (PAGA) framework to construct a coarse-grained graph. This facilitates the efficient computation of a structure-preserving geodesic distance matrix between meta-cells, where the distance metric integrates PAGA-inferred connectivity with the Euclidean distances between meta-cell centroids in a low-dimensional embedding (e.g., UMAP).

### Generalization to Multi-omics Features

Although we use the term “gene-centric” to align with established conventions in spatial transcriptomics, the GeneSPOT framework is inherently omics-agnostic. This framework represents the spatial distribution of any molecular features as a probability distribution over the tissue manifold. Since OT calculation measures only the effort required to reconfigure one distribution into another along the underlying transport geometry, GeneSPOT generalizes effectively to other spatially distributed molecular features. To implement this generalization, GeneSPOT employs specific strategies to address the distinct challenges presented by paired and unpaired multi-omics scenarios:

#### Paired Multi-omics Integration

When multiple omics layers are profiled on the same slice (e.g., spatial transcriptomics and metabolomics), the features are registered to a common spatial coordinate system. To integrate these layers, we first construct a unified spatial-aware cell graph by combining the diverse features into a holistic cellular representation (e.g., using SpatialGLUE^65^). Features from all modalities are then co-embedded into a unified Gene GCES. In this unified space, the proximity between a gene and a metabolite reflects similarity in their spatial deployment within the shared tissue context.

#### Unpaired Multi-omics Integration via Spatial Landmarks

Integrating unpaired datasets (e.g., transcriptomics and metabolomics from different samples) presents a unique challenge due to the absence of direct cell-to-cell correspondence. GeneSPOT addresses this challenge by leveraging shared spatial landmarks (i.e., alignable anatomical domains) as a conserved biological context to bridge disparate omics layers. The process begins by independently constructing a spatial-aware cell graph for each omics layer to capture its intrinsic tissue structure. To facilitate alignment, we employ a graph coarse-graining strategy to aggregate cells into meta-cells corresponding to the shared landmarks. By computing structure-preserving geodesic distances along the coarse-grained cell graph, GeneSPOT creates a comparable structural map for each omics layer. We then compute the average geodesic distance between landmarks across the different omics layers. Utilizing this averaged geodesic distance to calculate transport distances, GeneSPOT aligns features from separate experiments into a unified embedding space, enabling the discovery of cross-modal spatial associations without requiring physical overlap.

#### Quantifying Relational Spatial Dynamics Across Biological Contexts

GeneSPOT provides a quantitative and structured representation of relational spatial organization for all features, including measured genes and landmarks. Operationally, this organization comprises two complementary components: gene-to-landmark proximity, which measures anatomical anchoring, and gene-to-gene proximity, which measures spatial coupling among gene programs. By comparing context-specific GCESs, GeneSPOT quantifies relational spatial dynamics: changes in gene-to-landmark proximity reveal shifts in anatomical anchoring, whereas changes in gene-to-gene proximity characterize the reorganization of spatial coupling among genes. Proximity measures the spatial association between any two features and is calculated as the cosine similarity of their embedding vectors. For two features **F**_*i*_ and **F**_*j*_, the proximity is given by:

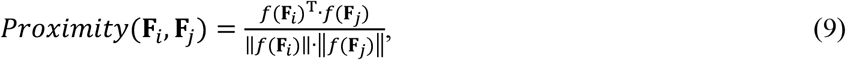

where ‖·‖ denotes the Euclidean L2 norm of a vector.

#### Gene-to-Landmark Proximity

To quantify a gene’s anatomical anchoring within each biological context, we calculate the proximity between the gene’s embedding and a landmark’s embedding within each context-specific GCES. Changes in this proximity across contexts indicate shifts in anatomical anchoring.

#### Gene-to-Gene Proximity

To quantify spatial coupling among genes within each biological context, we calculate the proximity between the embeddings of a gene pair within each context-specific GCES. Changes in this proximity across contexts characterize the reorganization of spatial coupling among genes, including convergence or divergence of their respective programs.

#### Gene Clustering

To identify modules of genes with similar spatial deployment patterns, we perform clustering on the gene embeddings within the GCES. First, a kNN graph is constructed on the embeddings by using “scanpy.pp.neighbors”. Then, we apply the Leiden community detection algorithm using the “scanpy.tl.leiden” to this graph to partition the genes into distinct gene clusters (GCs).

#### Gene set enrichment analysis

We performed over-representation analysis using Enrichr through the gseapy Python package, with Gene Ontology Biological Process 2025 as the reference gene-set library. Enrichment significance was assessed using one-sided Fisher’s exact tests with Benjamini–Hochberg correction, and terms with adjusted *P* < 0.05 were considered significant.

### Benchmarking and Validation Strategy

#### Evaluation Tasks

We evaluated prioritized features across five tasks using held-out test slices from multiple datasets:

- **Unsupervised PCA Separation Analysis:** This task evaluated whether prioritized genes capture sufficient biological variance to distinguish anatomical domains without supervision. Principal Component Analysis (PCA) was performed on a held-out test slice using only the expression matrix of genes prioritized from the training slices.
- **Supervised Domain Classification:** This task evaluated whether the selected genes carry sufficient discriminatory information to predict tissue architecture. A logistic regression model was trained on the training slices using the selected gene sets to predict anatomical domain labels in an unseen test slice.
- **Supervised Gene Expression Regression:** This task evaluated the capacity of selected genes to represent the broader transcriptional landscape. Using a leave-one-gene-out strategy, linear regression models were trained on the training slices to predict the expression of a held-out target gene based on the expression of selected feature genes in the unseen test slice.
- **Unsupervised Multi-omics Domain Identification:** This task evaluated cross-modal integration. We tested whether genes and metabolites prioritized in the training slice could reconstruct known tissue architecture in an unseen test slice using the SpatialGLUE algorithm.
- **Supervised Metabolite Regression:** Similar to gene regression, we trained models to predict the spatial abundance of specific metabolites (e.g., dopamine and 3-MT) on a test slice using gene features prioritized from a training slice.

#### Competing methods

We compared GeneSPOT against established comparison methods, baselines, and ablation models:

- **HVG (Highly Variable Gene) identification methods:** We implemented Seurat and CellRanger using the Scanpy package with default parameters to select genes based on expression variance.
- **SVG (Spatially Variable Gene) identification methods:** We utilized Moran’s I and Geary’s C (via the Squidpy package) and SpatialDE (via its official repository), all applied with default settings. Additionally, we benchmarked GeneSPOT against the recent gene-centric method STMiner (via its official repository) to evaluate feature prioritization.
- **Baseline:** To establish robust lower bounds for performance, we implemented several baselines. A Random Baseline was implemented by conducting 1,000 independent iterations of random feature selection from the entire feature panel. For multi-omics analysis, we implemented two correlation-based baselines: Pearson (domain-level), which ranks features based on their correlation with aggregated expression within annotated spatial domains, and Pearson (spot-level), which ranks features based on direct spot-to-spot correlation of raw abundances across all paired spots.
- **Ablation and Control Models:** To validate specific methodological components, we evaluated GeneSPOT-Pearson, a variant in which the structure-preserving OT distance was replaced with the standard Pearson correlation. We also assessed GeneSPOT-Perm, a control model in which spatial domain annotations were randomly shuffled prior to training to confirm that performance is driven by genuine spatial signals. Additionally, we implemented GeneSPOT-coords, an ablation variant that utilizes raw Euclidean coordinates to define the cell graph. This comparison demonstrated that resolving the underlying tissue structure is crucial for accurately modeling the functional landscape of genes.

#### Feature Selection Strategy

To prevent data leakage, all feature selection was performed exclusively on the training set, while evaluation was conducted solely on the held-out test set. For each evaluation task:

- **Unsupervised PCA Separation Analysis:** Gene selection was performed on the training set using the naive ranking criteria of each method—standardized variance for HVGs identification methods and spatial autocorrelation scores for SVGs identification methods. Specifically, the top set consisted of genes with the highest standardized variance or autocorrelation scores, while the bottom set included genes with the lowest scores. GeneSPOT prioritized genes based on the standard deviation (STD) of their proximities to spatial landmarks. The top set comprised genes with the highest STD, indicating high spatial specificity, whereas the bottom set included genes with the lowest STD, indicating uniform spatial distribution.
- **Supervised Domain Classification Task:** Similarly, gene selection was performed on the training set using the native ranking logic of each method to identify features for predicting anatomical labels. Specifically, for competing methods, the top set comprised genes with the highest standardized variance or autocorrelation scores, while the bottom set comprised genes with the lowest scores. For GeneSPOT, the top set consisted of genes with the highest STD of their proximities to spatial landmarks, while the bottom set consisted of those with the lowest STD.
- **Supervised Gene Expression Regression Task:** Genes were selected based on their relative proximity to the held-out target gene. For the HVGs and SVGs methods, we calculated the pairwise differences in their naive statistical scores. The top set comprised genes with the smallest differences in scores relative to the target gene, while the bottom set comprised genes with the largest differences. For GeneSPOT, we prioritized genes based on their gene-to-gene proximity to the target gene in the GCES. Specifically, the top set comprised genes with the highest gene-to-gene proximity to the target gene, while the bottom set comprised those with the lowest proximity.
- **Unsupervised Multi-omics Domain Identification Task:** We applied a feature selection strategy consistent with that used in the supervised domain classification task independently to both gene and metabolite modalities, ensuring that each method employed its native metric to identify the most informative features.
- **Supervised Metabolite Regression Task:** We employed a strategy analogous to the gene regression task to identify gene features for predicting target metabolites (specifically dopamine and 3-MT), utilizing cross-modal proximity for GeneSPOT and statistical score differences for baseline methods.

#### Evaluation Metrics

To ensure the robustness of our results, we evaluated performance across a wide range of feature set sizes. We employed specific metrics for each task to quantify performance.

- **Unsupervised PCA Separation Analysis:** The capacity of the prioritized genes to distinguish tissue structures without supervision was quantified using the number of significantly separated domain pairs. Specifically, the distribution of principal component scores was compared between every possible pair of distinct anatomical domains. For each pair, the statistical significance of the separation was determined using an unpaired two-sided t-test. The domain-separation score was defined as the number of anatomical-domain pairs with different PC-score distributions at a nominal two-sided *P*<0.05. A higher count indicates that the selected genes effectively capture the intrinsic structural variance of the tissue.
- **Supervised Domain Classification Task:** The ability of the prioritized features to correctly assign cells to ground-truth anatomical domains was quantified using Accuracy, F1-Score, and the Area Under the Receiver Operating Characteristic Curve (ROC-AUC). These evaluation metrics can be formulated as follows:

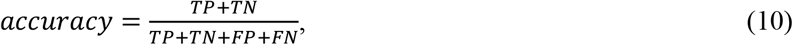

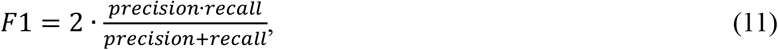

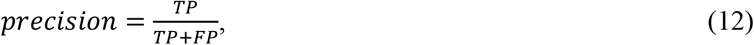

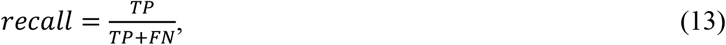

where *TP, TN, FP*, and *FN* are the count of true positives, true negatives, false positives, and false negatives, respectively.
- **Supervised Gene Expression Regression Task:** The capacity of the selected gene set to represent the overall transcriptional landscape was evaluated using Pearson Correlation (*r*), R-squared (*R*^2^), and Explained Variance (*EV*), which are formulated as:

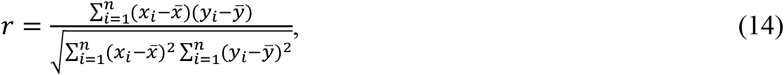

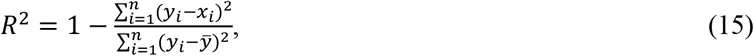

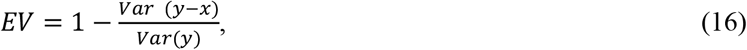

where *x* and *y* represent the predicted value and ground truth, and 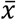and 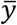 indicate their mean values, *Var*(·) is the variance of the values.
- **Unsupervised Multi-omics Domain Identification Task:** The effectiveness of integrated multi-omics features in delineating spatial domains was evaluated by comparing the identified clusters with anatomical annotations. The concordance was quantified using the Adjusted Rand Index (ARI). This metric can be formulated as follows:

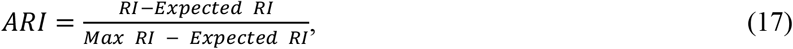

where the *RI* is the Rand Index, defined as:

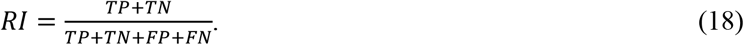
- **Supervised Metabolite Distribution Regression Task:** The ability of prioritized genes to predict the spatial distribution of specific metabolites was quantified using Pearson Correlation (*r*), Cosine Similarity (*θ*), and Explained Variance (*EV*). The Cosine Similarity can be formulated as follows:

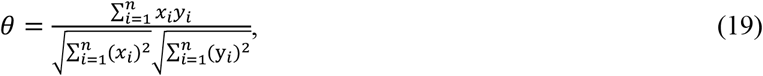

where *x* and *y* represent the predicted value and ground truth metabolite abundance.

## Statistics & Reproducibility

This study is a computational reanalysis of previously published, publicly available spatial omics datasets described in Supplementary Notes 1; no new wet-lab experiments were performed. No statistical method was used to predetermine sample size. Dataset sizes were determined by the original data-generating studies, and all available cells, genes and tissue slices meeting the predefined quality-control criteria in Supplementary Notes 1 were included; the sample sizes therefore reflect the complete eligible data available for each analysis. Beyond the stated quality-control and feature-filtering criteria, no data were excluded. Because the study reanalyzes observational datasets rather than a controlled experiment, the analyses were not randomized, and investigators were not blinded to sample identity. Reproducibility and robustness were assessed using held-out test slices, sensitivity analyses across feature-set sizes and control analyses including random feature selection and GeneSPOT-Perm, as described in the relevant Methods sections and figure legends.

For pairwise statistical comparisons, two-sided unpaired Student’s t-tests were used for PCA domain-separation analyses (Figure 2F and Supplementary Figure 9), and two-sided Spearman correlation tests were used for gene-to-landmark and gene-to-gene proximity analyses (Figure 5). These analyses report nominal *P* values at *P* < 0.05; no adjustment for multiple comparisons was applied. Gene set enrichment analyses instead used Benjamini–Hochberg false-discovery-rate-adjusted *P* values (adjusted *P*<0.05), as implemented in gseapy/Enrichr. All quantitative benchmarking comparisons were performed on held-out test data not used for feature selection or model training.

## Implementation details

This study was implemented using the PyTorch platform. The models were trained using the Adam optimizer, with the learning rate set as 1e-3. Model training was conducted for a total of 200 epochs, and batch sizes were configured as 128. All experiments were conducted on a server running Ubuntu, equipped with an Intel(R) Xeon(R) Gold 5418Y CPU (2.00 GHz), 503 GB of RAM, and an NVIDIA A800 GPU (80 GB).

## Data availability

All data analyzed during this study are publicly available. The STARmap mPFC dataset is available from https://github.com/zhengli09/BASS-Analysis/tree/master/data. The BaristaSeq Visual cortex data is available from https://spacetx.github.io/data.html. The mouse hypothalamus MERFISH dataset is available via Dryad from https://doi.org/10.5061/dryad.8t8s248 (Ref^98^). The SMA dataset is available via figshare at https://doi.org/10.17044/scilifelab.22770161 (Ref^99^). The 10x breast cancer dataset is available from https://www.10xgenomics.com/products/xenium-in-situ/preview-dataset-human-breast. The spatially resolved transcriptomics of mouse kidney is available from Gene Expression Omnibus under accession number GSE266244. The spatially resolved metabolomics of mouse kidney is available from https://metaspace2020.org/datasets?q=2023-12-19_13h02m39s. The Stereo-seq mouse embryo development dataset used in this study is available from the CNGB database under accession code CNP0001543. The single-cell spatial transcriptomics atlas of the aging mouse brain is available via Zenodo from https://doi.org/10.5281/zenodo.13883177 (Ref^100^).

## Code availability

The Python implementation of the GeneSPOT framework is publicly available on GitHub at https://github.com/YSTLab/GeneSPOT.

